# Dynamic decoupling of biomass and lipid biosynthesis by autonomously regulated switch

**DOI:** 10.1101/337790

**Authors:** Suvi Santala, Elena Efimova, Ville Santala

## Abstract

For improving the microbial production of fuels and chemicals, gene knock-outs and overexpression are routinely applied to intensify the carbon flow from substrate to product. However, their possibilities in dynamic control of the flux between the biomass and product synthesis are limited, whereas dynamic metabolic switches can be used for optimizing the distribution of carbon and resources. The production of single cell oils is especially challenging, as the synthesis is strongly regulated, competes directly with biomass, and requires defined conditions, such as nitrogen limitation. Here, we engineered a metabolic switch for redirecting carbon flow from biomass to wax ester production in *Acinetobacter baylyi* ADP1 using acetate as a carbon source. Isocitrate lyase, an essential enzyme for growth on acetate, was expressed under an arabinose inducible promoter. The autonomous downregulation of the expression is based on the gradual oxidation of the arabinose inducer by a glucose dehydrogenase *gcd*. The depletion of the inducer, occurring simultaneously to acetate consumption, switches the cells from a biomass mode to a lipid synthesis mode, enabling the efficient channelling of carbon to wax esters in a simple batch culture. In the engineered strain, the yield and titer of wax esters were improved by 3.8 and 3.1 folds, respectively, over the control strain. In addition, the engineered strain accumulated wax esters 19% of cell dry weight, being the highest reported among microbes. The study provides important insights into the dynamic engineering of the biomass-dependent synthesis pathways for the improved production of biocompounds from low-cost, sustainable substrates.

**Significance statement:** In the biological production, one of the greatest challenges is to find ways for optimal distribution of resources between cell growth, maintenance, and product synthesis. Robust and reliable circuits are required to allow autonomous switching of cells from biomass mode to lipid synthesis mode. Dynamic production of single cell oils such as triacylglycerols and wax esters is especially challenging due to the strong regulation. We present a dynamic genetic circuit based on conditional knockdown of a glyoxylate shunt enzyme, which is essential for cell growth. By gradual repression of the gene, the cells autonomously switch from biomass mode to product synthesis mode. We demonstrate the functionality of the circuit by using bacterium *Acinetobacter baylyi* ADP1 for the production of long chain alkyl esters, namely wax esters, with titer and yield improved by over 3-fold using acetate as the carbon source.

## 1. Background

Metabolic engineering and synthetic biology provide powerful means for the bio-based production of a variety of chemicals and other commodities by engineered microbes. Compared to conventional chemical synthesis, the superiority of biological production systems lies in the possibility to synthesize both the catalyst (i.e. the cell factory) and the product itself from very simple chemical compounds, such as sugars or organic acids. However, the challenge is to develop a system, which optimally distributes the resources and carbon flux between building up the catalyst and operating the catalyst for the actual production; extensive cell growth takes resources from the product synthesis, whereas too excessive flux towards product synthesis may result in reduced growth, insufficient cofactor regeneration, low enzyme expression levels, and eventually poor titers. To address the challenges related to the optimal distribution of cellular resources, a number of dynamic circuit designs targeting the central pathway nodes have been recently developed (1). While growth-associated genes responsible for central carbon metabolism cannot be directly deleted, various strategies for decoupling growth and product synthesis have been introduced; Soma et al. constructed a metabolic toggle switch for conditional knockout of citrate synthase *gltA,* an enzyme required for functional tricarboxylic acid (TCA) cycle (2). The switch allowed an induced shift of carbon flow from TCA cycle to synthetic isopropanol pathway. More recently, the system was further improved by introducing a sensor-regulator system responsive to a defined cell density (3). Solomon et al. introduced a dynamic approach to controlling the glycolytic flux; antisense RNA technology and an inverting gene circuit were employed for inhibiting the activity of glucokinase (Glk), resulting in a controlled growth rate and a reduced production of acetate (4). Brockman and Prather introduced another example of a dynamic regulation system, where they developed a circuit for dynamic knockdown of phosphofructokinase-1 (Pfk-1), the enzyme responsible for the key step in the glycolytic pathway regulating glucose-6-phosphate flux. By the temporal control of Pfk-1 degradation, glucose-6-phosphate could be efficiently directed to heterologous *myo*-inositol synthesis pathway instead of biomass production (5). In a previous work, Doong et al. further improved the system with a *myo*-inositol responsive dynamic sensor that regulated the downstream enzymes of the pathway in converting *myo*-inositol to glucarate. The introduced systems represent highly elegant examples of advanced metabolic control, but the complex circuit designs can be prone to destabilization in prolonged cultivations and function unexpectedly in scaled-up processes (6). Thus, reliable metabolic control systems with simple and robust operation would be also desired.

Microbial storage compounds, such as polyhydroxyalkanoates (PHA), triacylglycerols (TAG) and wax esters (WE), are industrially relevant and desirable molecules due to their vegetable oil like properties and broad applicability in e.g. fuel, nutritional, cosmetic, and pharmaceutical industries. However, the production of the long carbon chain products derived from fatty acyl-CoA requires significant energy investments from the cell. In addition, the synthesis of storage compounds directly competes with biomass production and is strongly regulated, growth-phase dependent, and requires high amounts of cofactors and excess carbon along with limitation on other nutrients, such as nitrogen (7, 8). While strategies for dynamically regulated production of free fatty acids (FFA) and FA derived products have been introduced (9, 10), means for overcoming the challenges of storage lipid synthesis regulation are still lacking. Therefore, the production processes are conventionally improved by non-specific means, such as bioprocess optimization or in conditions with a defined carbon-nitrogen ratio (11). As an example of a more advanced approach, Xu et al. established a semi-continuous culture system with model-aided bioprocess optimization and cell recycling, which allowed efficient TAG production and high productivities even with dilute acetate feed (12). While also metabolic engineering strategies have been employed for improving TAG (13-16) and WE synthesis (17, 18) in microbes, efficient overproduction of acetyl-CoA coupled with dynamic resource distribution between biomass and storage lipid synthesis remains a challenge.

When microbes are cultivated on non-glycolytic substrates, such as acetate or fatty acids, complete TCA cycle yielding carbon dioxide cannot be utilized for anabolic processes. In those circumstances, the cells have to rely on an alternative route, namely glyoxylate cycle, which bypasses the two decarboxylation steps of TCA cycle. The key enzyme in the glyoxylate cycle is isocitrate lyase (AceA), responsible for converting isocitrate to glyoxylate and succinate. It was previously demonstrated that a knock-out of the isocitrate lyase in *Pseudomonas putida* improved the production of PHAs when grown on gluconate, apparently for providing surplus acetyl-CoA for the PHA synthesis (19). Some bacteria also exhibit an alternative pathway for glycolysis, such as the modified Entner-Doudoroff pathway of certain *Acinetobacter* strains (20, 21). An interesting feature of the glycolysis of *Acinetobacter* is the oxidation of glucose to gluconate prior the transport to cells. In the absence of glucose, the first enzyme of the pathway, glucose dehydrogenase Gcd, can unselectively oxidize other sugar compounds present in the medium, such as pentoses, without the capability to utilize them as a carbon source (22-24). Importantly, this feature does not interfere with the utilization of non-glycolytic carbon sources, such as organic acids, and can be considered being ‘orthogonal’ to the substrate utilization. Thus, this feature could be exploited in the regulation of metabolic pathways in the cells.

Here, we construct a dynamic switch for autonomous shifting of cells from the biomass mode to the storage lipid synthesis mode by introducing a circuit for a conditional elimination of the glyoxylate cycle, which is the essential bypass for cells growing on acetate, and the key control node in lipid biosynthesis pathway. We demonstrate the functionality of the switch by the improved (both yield and titer) production of wax esters using *A. baylyi* ADP1 as a host.

## 2. Material and Methods

### Strains and molecular work

*A. baylyi ADP1* (DSM 24193, Deutsche Sammlung von Microorganismen und Zellkulturen, Germany) was used in the study. The single gene knock-out strain of *A. baylyi* ADP1δ*aceA*∷ *tdk*/*Kan*^*r*^(ACIAD1084 deleted) was kindly provided by Veronique de Berardinis (Genoscope, France). In the single gene knock-out mutant, the gene in question is replaced with a gene cassette containing a kanamycin resistance gene (Kan^r^) (25). For cloning and plasmid amplification, *Escherichia coli* XL1-Blue (Stratagene, USA) was used. The reagents and primers for molecular work were purchased from ThermoFisher Scientific (USA), unless stated otherwise. The primers used in the study are presented in Table 1. The complementation of *aceA* in the knock-out strain was carried out by constructing a gene cassette with the *aceA* gene under an arabinose inducible promoter AraC-pBAD; a previously described gene cassette (16, 26) was used as the scaffold for the construction. The arabinose promoter and AraC repressor, designated as ara, were ampli?ed from pBAV1C-ara-LuxCDE plasmid (18) using primers VS10_09 and VS10_10 and inserted to the plasmid iluxAB/pIX (26) using restriction sites MfeI and NdeI. The gene *aceA* (ACIAD1084) was amplified from the genome of ADP1 wt with primers SS17_08 and SS17_09 and cloned to the ara-iluxAB/pIX using restriction sites NdeI and XhoI. Transformations of ADP1 were carried out as described previously (16). The transformed cells were selected on LA plates containing 25 µg/ml chloramphenicol. ADP1 strain with a *poxB* deletion, *A. baylyi* ADP1δ*poxB*∷ *Cm*^*r*^, was used as the reference strain designated as ADP1 wt. All genetic modifications were confirmed with PCR and sequencing.

**Table 1.**
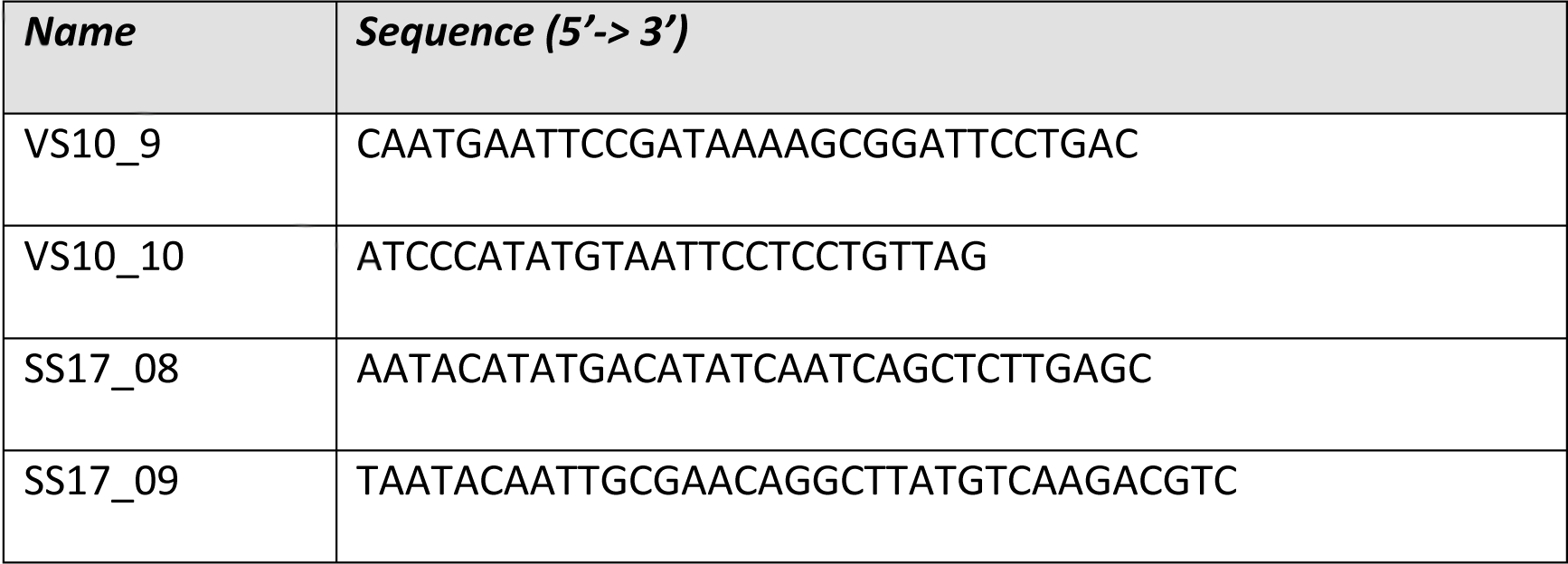
List of primers used in the study.

### Cultivations

The components for culture mediums were purchased from Sigma (USA). The strains were cultivated in modified MA/9 minimal salts medium (17) at 25 °C and 300 rpm unless stated otherwise. The medium was supplemented with 25-100 mM Na-acetate or 250 mM glucose, 0.1 % casein amino acids (w/v), and 0-1.0 % L(+)arabinose when appropriate.

The cultivations for optimizing the arabinose concentration for ADP1-ara-aceA growth were carried out using Tecan Spark^®^ (Tecan, Switzerland) microplate reader in 25 °C for 21 hours with two replicate wells for each strain and arabinose concentration. The medium was supplemented with 25 mM Na-acetate and 0.1 % casein amino acids, and arabinose (concentrations 0; 0.05; 0.1; 0.2; 0.5 or 1.0 %) was used for the induction of *aceA*. Mediums without acetate supplementation was used as the control medium to determine the growth on casein amino acids. The strain ADP1 wt was used as the positive control, whereas the knock-out strains ADP1 δ*aceA*∷ *Tdk/kan*^*r*^and *A. baylyi* ADP1δ*aceA*∷ *tdk*/*Kan*^*r*^δ*poxB*∷ *Cm* were used as negative controls for growth on acetate. For semi-quantitative determination of the WE production of ADP1-ara-aceA, the cells were cultivated in two parallel 5 ml cultures in the same medium except with 50 mM Na-acetate in 5 ml tubes for 30 hours.

For the quantitative determination of the WEs with NMR, the strain ADP1 wt and ADP1-ara-aceA were cultured in total 600 ml of medium supplemented with 50 mM Na-acetate and 0.5 % (~30 mM) arabinose distributed in 12 Erlenmeyer flasks. HPLC samples were taken every 1-5 hours and two parallel 40 ml samples were taken in five different time-points to quantitatively determine WE (by NMR) and biomass production (as cell dry weight).

### Lipid and end-metabolite analyses

The amount of total lipids and WEs were estimated by TLC or quantified by NMR. For TLC, equal volumes of samples (3 ml) from different cultures were taken and the lipids were extracted using ‘miniscale’ chloroform-methanol extraction as described previously (17). Thirty µl of the chloroform phase was applied on 20 × 10 cm Silica Gel 60 F_254_ HPTLC glass plates with 2.5 × 10 cm concentrating zone (Merck, USA). Mobile phase used was n-hexane: diethyl ether: acetic acid 90: 15: 1 and iodine was used for visualization. Jojoba oil was used as the standard for WEs. For comparative evaluation of the intensities of the WE bands on TLC, the Gel analysis method of ImageJ software (rsb.info.nih.gov/ij/index.html) was applied as described in the ImageJ documentation.

For NMR analyses, the 40-ml biomass samples were freeze-dried and the cell dry weight (CDW) was determined gravimetrically. The lipid extraction and the quantitative ^1^H NMR analysis of WEs was carried out as described earlier (26). The amount of total lipids was determined gravimetrically. The areas of the peaks in the NMR spectrum are directly proportional to the molar concentration of each functional group, yielding specific concentration for WEs in total biomass. The concentration of WEs was calculated from the integrated signal at 4.05 ppm which is characteristic for protons of α-alkoxy-methylene group of esters (–CH2-COO-CH2–). For the calculation of the WE titer in grams per liter, an average molar mass of 506 g/mol was used (17).

The glucose, acetate, and arabinose concentrations were determined by LC-20AC prominence liquid chromatograph (Shimadzu, USA) equipped with RID-10A refractive index detector, DGU-20A5 prominence degasser, CBM-20A prominence communications bus module, SIL-20AC prominence autosampler, and Shodex SUGAR SH1011 (Showa Denko KK, Japan) as described previously (16).

## 3. Results

In order to investigate the effect of the knock-out of isocitrate lyase AceA on the growth and WE production in *A. baylyi* ADP1, we employed a knock-out mutant strain *A. baylyi* ADP1 δ*aceA*∷ Tdk/kanr (25) for preliminary test cultivations. We observed that when grown on glucose, the cells grow more slowly, but produce WE titers comparable to those of the wild type (wt) strain; after 48 hours of cultivation, the wild type had produced 470±150 mg/l WEs compared to 460±40 mg/l WEs produced by the knock-out strain. In opposite to the wt strain, however, the mutant strain did not exhibit growth on minimal medium supplied with acetate as the sole carbon source. This is due to the lack of route for acetyl-CoA to be directed in biosynthetic pathways via malate. Thus, as acetyl-CoA represents the key precursor in both the biomass production through the glyoxylate shunt and the wax ester biosynthesis, we hypothesized that by dynamically regulating the isocitrate lyase, the state of the cells could be switched between biomass and lipid synthesis modes (Fig 1). In order to make the shift dynamic, we introduced an approach for autonomous regulation of the isocitrate lyase AceA; by expressing the enzyme under an arabinose-inducible promoter AraC-pBAD, the induction is gradually repressed due to the depletion of arabinose by the glucose dehydrogenase activity of ADP1. The arabinose inducible promoter has been previously shown to function in *A. baylyi* ADP1 (18, 27). In order to establish a system with maximal linearity and controllability, we constructed a gene cassette for genomic expression of *aceA* (Fig 1c, d). Exploiting the natural transformation machinery of ADP1, the gene cassette was integrated in the genome to replace a gene *poxB* (ACIAD3381), which has been previously shown to be a neutral target site in terms of growth and wax ester production (26, 28). The resulting strain *A. baylyi* ADP1δ*aceA*∷ *tdk*/*Kan*^*r*^δ*poxB*∷ *araC*-pBAD-*aceA*-*Cm*^*r*^was designated as ADP1-ara-aceA. The strain *A. baylyi* ADP1δ*poxB*∷ *Cm*^*r*^was used as the reference strain, from now on designated as the ADP1 wt.

**Figure 1.**
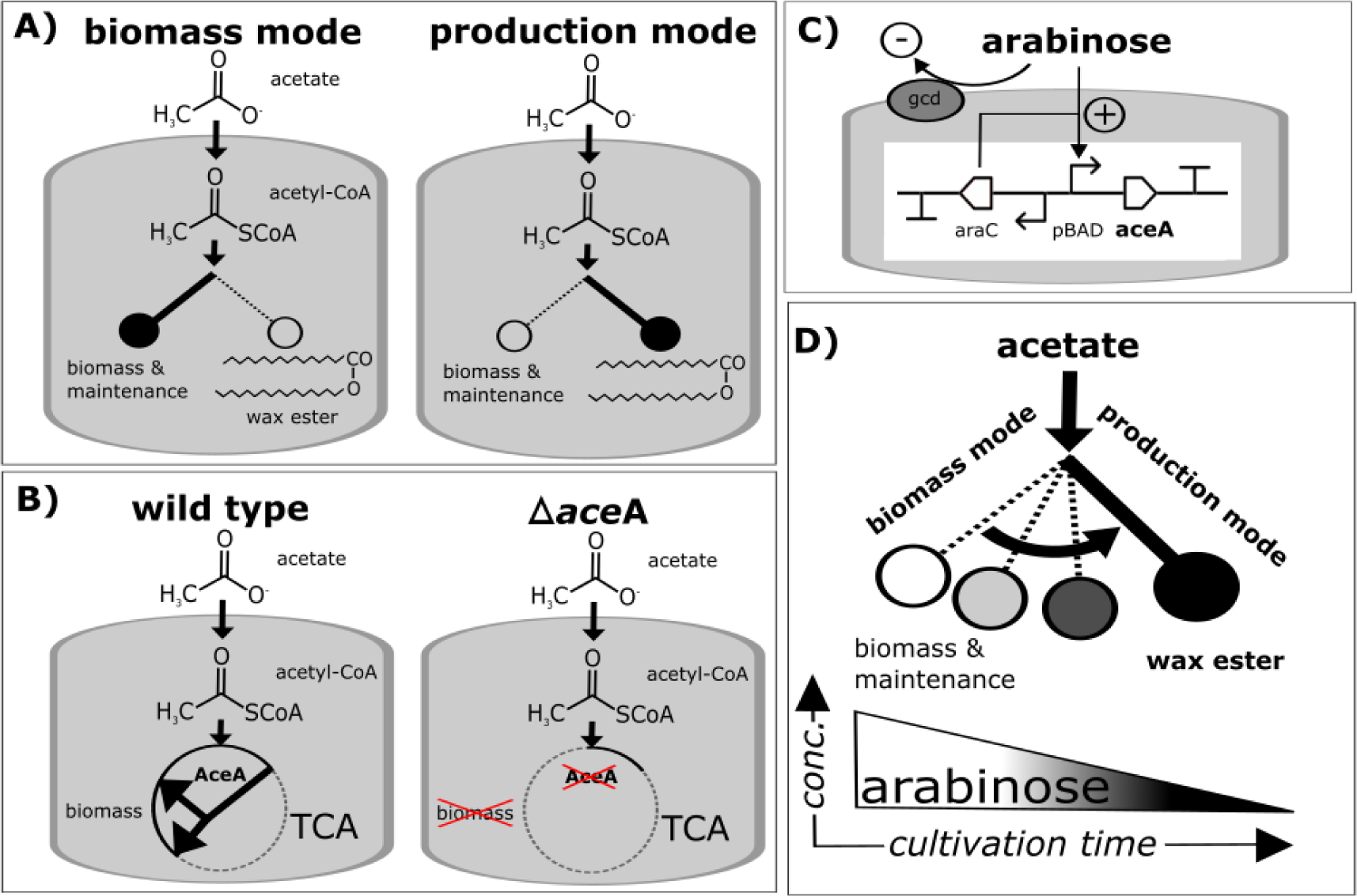
An autonomously regulated switch for shifting the cells from a biomass to a product synthesis mode. A) Wax ester synthesis strongly competes with biomass production; In the wild type cells, efficient wax ester synthesis is triggered only in defined conditions (i.e. in conditions with nitrogen starvation and excess carbon). In normal growth conditions, most of the carbon is directed to biomass production and cell maintenance. B) The ADP1 wild type strain utilizes acetate for biomass production through a glyoxylate shunt in citric acid cycle. Isocitrate lyase (AceA) is the key enzyme in the glyoxylate shunt and thus essential for growth on acetate. Strain lacking this enzyme is unable to grow on acetate as the sole carbon source. C) The genetic construct for dynamic regulation of *aceA* expression and growth. In the construct, *aceA* is placed under the arabinose-inducible promoter AraC-pBAD. When cultured on acetate, the arabinose used as the inducer is slowly oxidized by the native glucose dehydrogenase Gcd of ADP1, gradually repressing the expression of AceA. Arabinose oxidation does not serve as the carbon source or interfere with the acetate utilization, thus being orthogonal to the circuit function. D) Along with the inducer depletion (arabinose oxidation), the cells gradually shift from the biomass mode to the lipid synthesis mode; the less there is arabinose left in the culture, a higher proportion of the carbon flux is directed to the product. The amount of biomass can be simply regulated by adjusting the initial arabinose concentration.

We investigated the functionality of the complementation in the strain ADP1-ara-aceA in minimal medium supplemented with 50 mM acetate (Figure 2). Arabinose (at concentrations 0; 0.1 and 1.0 %) was added to the cultures, and the cultivations were continued for 68 hours. Cells did not grow or consume acetate in the absence of arabinose, indicating sufficiently tight regulation of the arabinose promoter. In the cultures with small amount of arabinose (0.1 %), the cells stopped growing after reaching an optical density (OD) of 0.3 and consumed only 5 mM acetate, whereas with 1.0 % arabinose the cells reached OD ~3.5 along with complete consumption of acetate. The ADP1 wt grew to slightly lower biomass (OD ~2.4) and consumed all the acetate; arabinose supplementation had no effect on the ADP1 wt growth. After 68 hours of cultivation, approximately 85 % of the 1% arabinose had been oxidized. The control knockout strains ADP1δ*aceA∷ tdk/Kan*^*r*^ and ADP1*δaceA∷ tdk/Kan*^*r*^ *δpoxB∷ Cm*^*r*^did not exhibit growth nor acetate consumption with or without the presence of arabinose (OD 0 at 0-68 h). We also confirmed, that the AceA expression is repressed due to arabinose oxidation, i.e. the conversion of arabinose to non-inducive form, arabino-lactone and further to arabonate (Figure S1).

**Figure 2.**
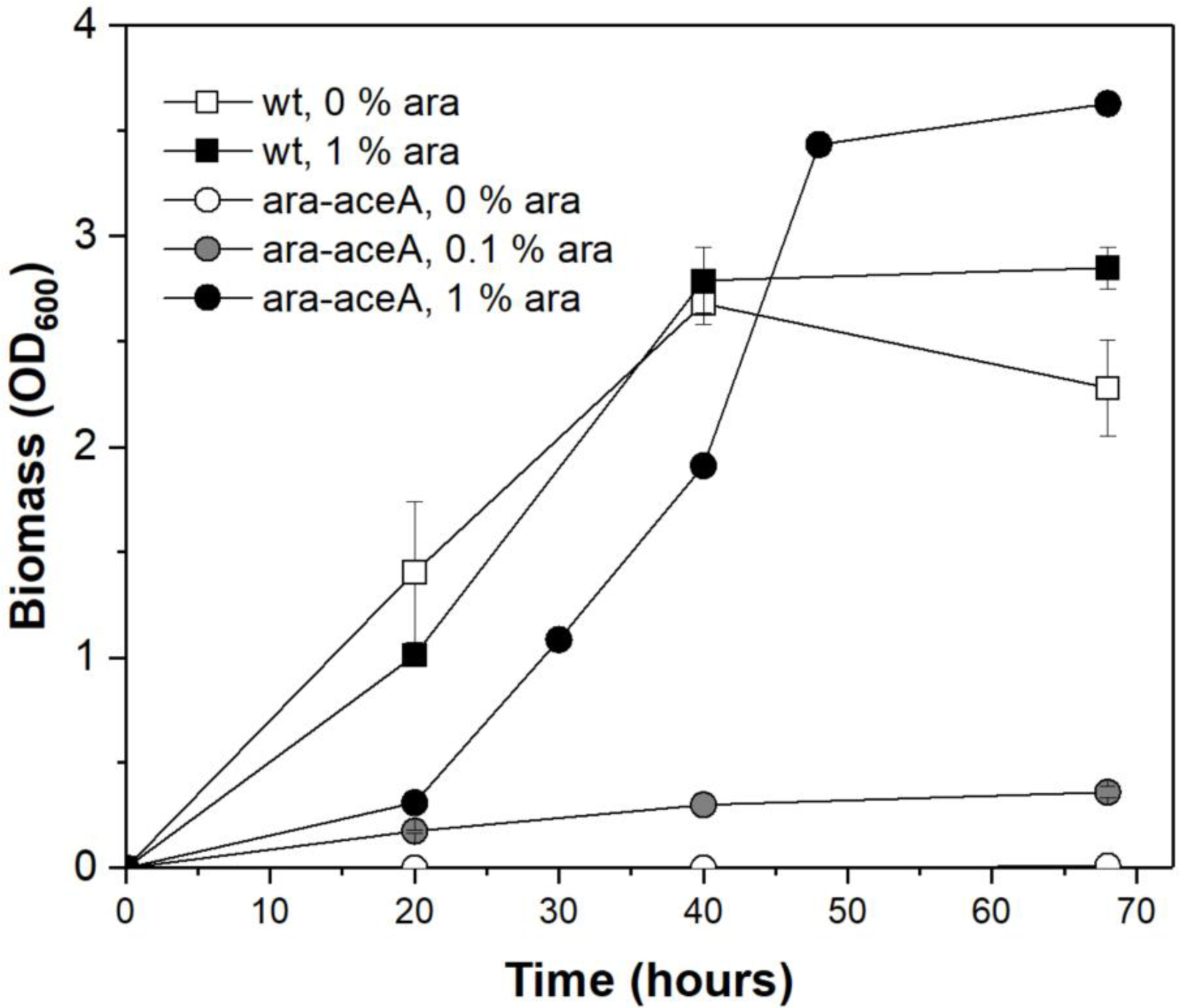
Growth of ADP1 wt and the engineered strain ADP1-ara-aceA in minimal medium. The cells were cultured for 68 hours at 25 °C in MSM supplemented with 50 mM Na-acetate as the sole carbon source. Arabinose concentrations of 0; 0.1 and 1.0% (for ADP1 wt 0 and 1%) were used for the induction. Optical densities representing biomasses are presented as an average of two individual replicates.

In order to find the optimal arabinose concentration in terms of both biomass and wax ester production, the strain ADP1-ara-aceA was cultivated in several different arabinose concentrations in minimal salts medium supplemented with acetate (Figure 3) for 21 hours. ADP1 wt was cultured as the reference strain. Casein amino acids (0.1%) were added to the culture in order to promote the growth and to prevent nitrogen limitation. As indicated by the previous growth experiment, we found that 1% arabinose was sufficient to allow the engineered strain to reach the same biomass as ADP1 wt, albeit the cells grew slower. Within the concentration range 0 – 0.2%, only small differences in growth pattern or biomass production were observed. The slight increase in OD of uninduced cells is due to the utilization of the casein amino acids present in the growth medium; a same amount of biomass is achieved without acetate supplementation with the wild type strain and the knock-out strain ADP1*δaceA∷ tdk/Kan*^*r*^*δpoxB∷ Cm*^*r*^with both acetate and casein amino acid supplementation (data not shown). For ADP1 wt, all the growth curves were similar regardless of the arabinose concentration used (data not shown).

**Figure 3.**
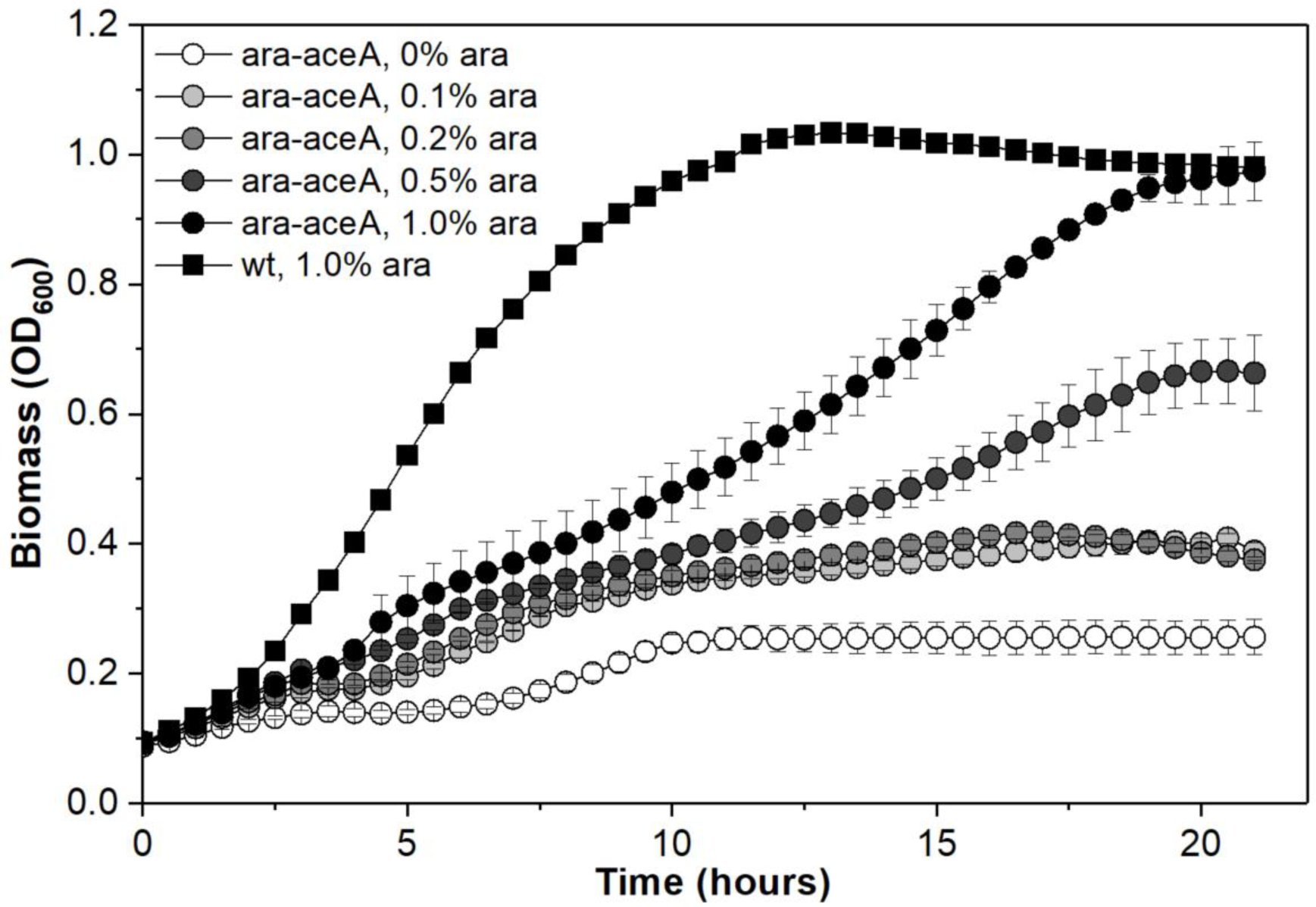
Growth of ADP1-ara-aceA and ADP1 wt with different arabinose concentrations. The cells were cultured in MA/9 medium supplemented with 25 mM acetate, 0.1% cas.amino acids, and arabinose (0-1%) at 25 °C in micro well-plates for 21 hours. The optical densities are presented as the average of three different replicate wells. For ADP1 wt, only the growth curve of the culture containing 1% arabinose is shown.

Next, we determined which initial arabinose concentration most optimally distributes the carbon between the biomass and the WE production. The strain ADP1-ara-aceA was cultivated in 5 ml minimal salts medium supplemented with 0.1 % casamino acids and 50 mM acetate with different arabinose concentrations (0; 0.1; 0.2; 0.5% and 1.0%) for 30 hours, after which biomass and WE production were determined. A clear correlation between the arabinose concentration and biomass production (OD600) was detected (Figure 4). Without arabinose supplementation, the cells grew to an OD of approximately 0.8, which is due to the utilization of casamino acids (the same OD was obtained with ADP1*δaceA∷ tdk/Kan*^*r*^*δpoxB∷ Cm*^*r*^). A semi-quantitative lipid analyses based on thin layer chromatography (TLC) was carried out to compare the amount of WEs produced (Figure 5B). For all the cultures, the same sample volume was taken for the analysis, thus representing the titer of WEs produced. Based on image analysis, the intensity of the band representing the wax ester titer increases along with the biomass concentration. However, only a slight difference was observed between the bands of the 0.5% and 1.0% cultures, suggesting that in the culture which contains saturating amount of arabinose the growth corresponds to that of the wild-type strain. When the intensities were divided with the optical densities, the highest amounts of WEs (per cell) were produced in the cultures with 0.2% and 0.5% arabinose. Thus, considering both the volumetric titer and the yield of WEs per biomass, the arabinose concentration of 0.5% was found to be optimal in terms of distributing the carbon and cellular resources between biomass and WEs.

**Figure 4.**
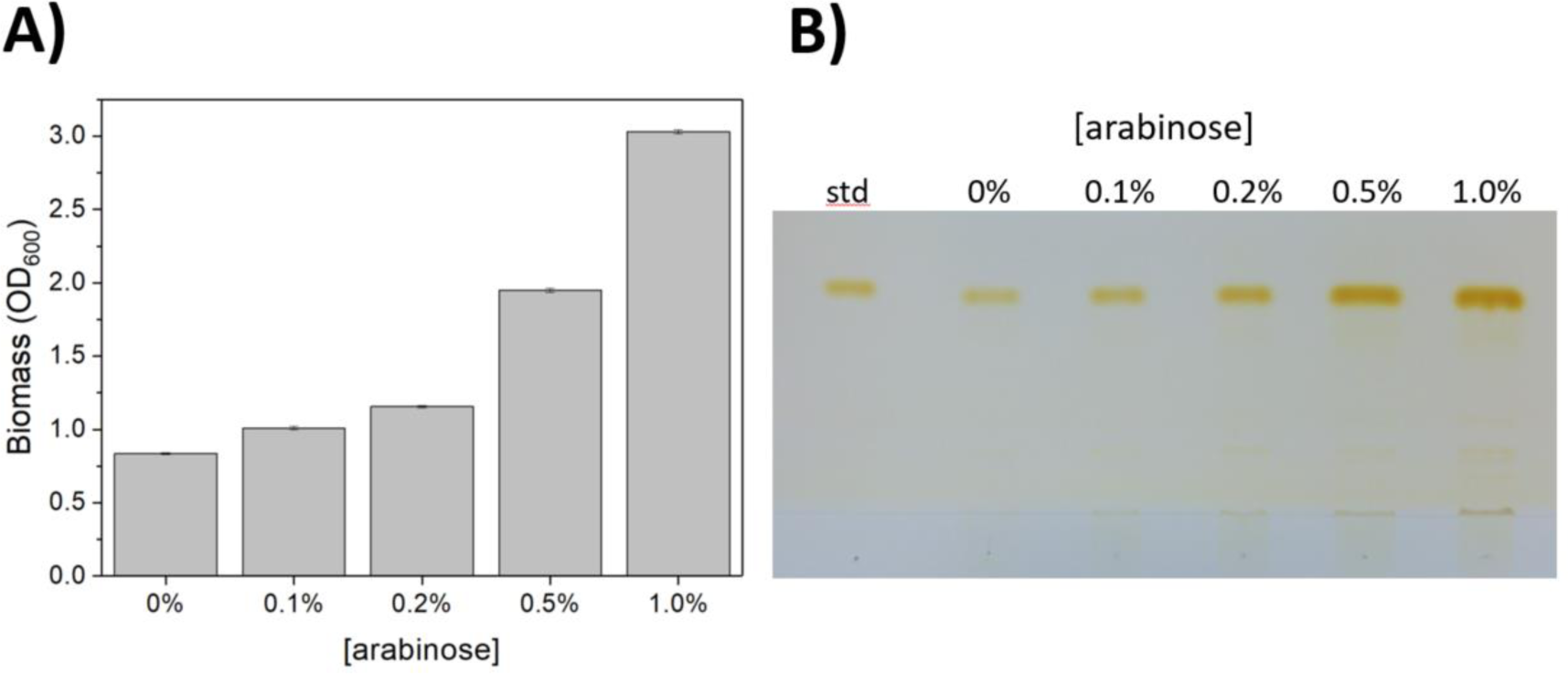
Biomass and wax ester production by ADP1-ara-aceA induced with different arabinose concentrations. ADP1-ara-aceA was cultured in MA/9 supplemented with 50 mM acetate, 0.1% casam, and arabinose (0, 0.1, 0.2, 0.5, or 1.0%) for 30 hours. A) The amount of the produced biomass (at the end-point) was determined by optical density (600 nm) measurement, presented as an average of two individual replica cultures. B) Wax ester production in the cultures with different arabinose concentrations were determined by thin layer chromatography analysis, for which 3 ml samples for taken from each culture. Jojoba oil was used as the standard.

**Figure 5.**
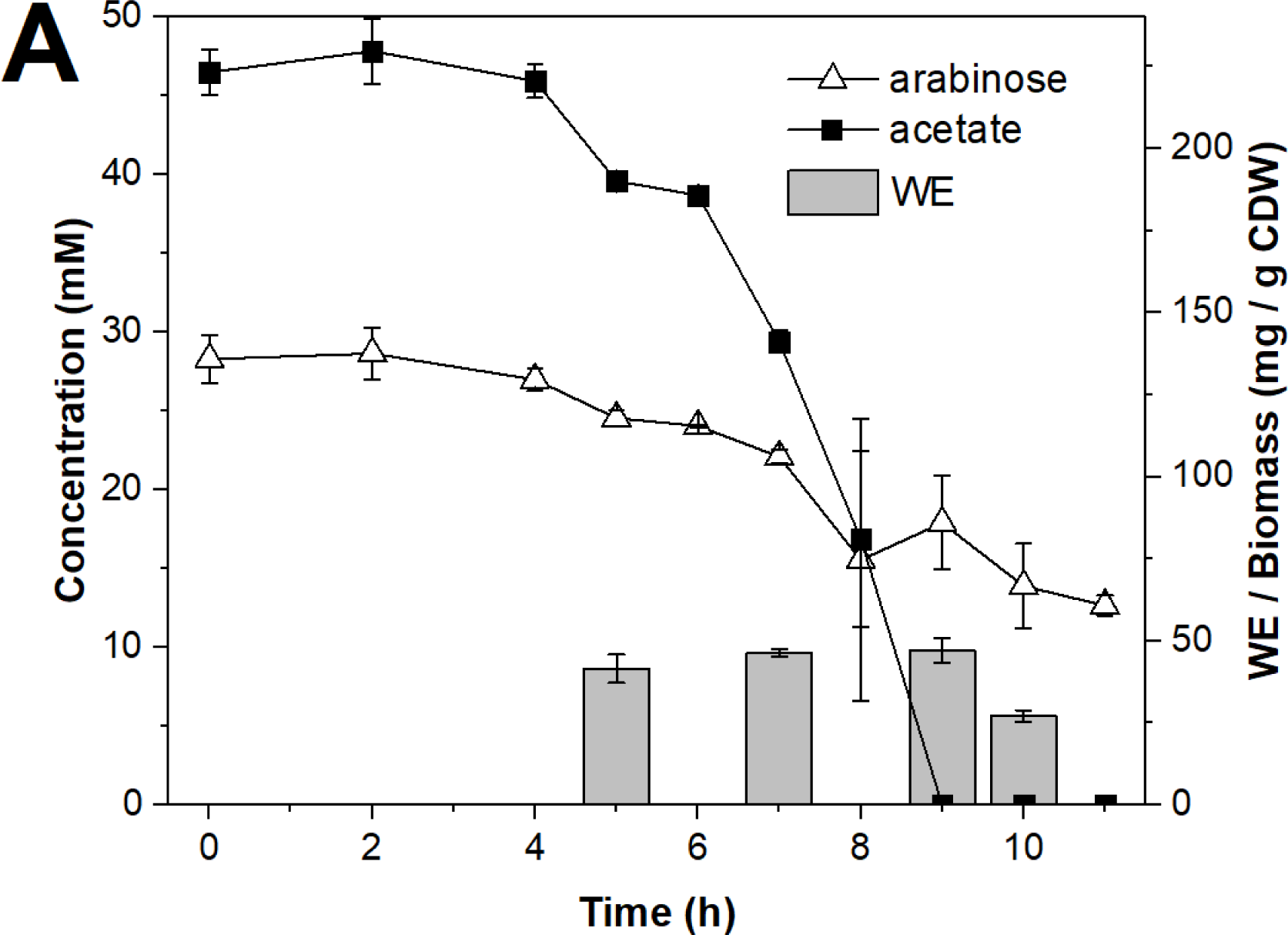
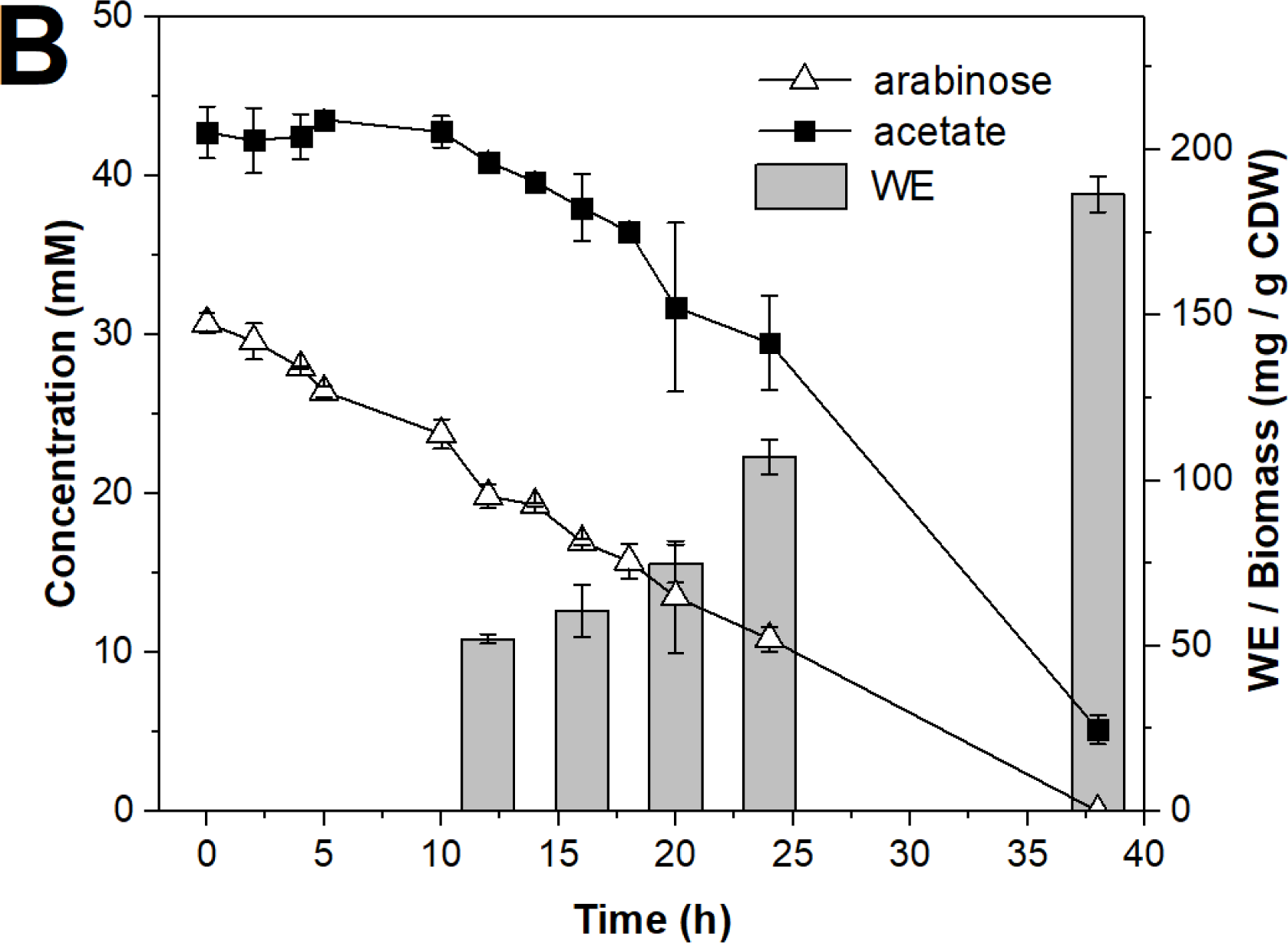
Acetate utilization, arabinose oxidation, and the accumulation of wax esters (WE) in the batch cultures of A) ADP1 wt and B) ADP1-ara-aceA. The cells were cultivated in MA/9 medium supplemented with 50 mM acetate, 0.1% cas. amino acids, and 0.5% arabinose for 12 h (ADP1 wt) or 38 h (ADP1-ara-aceA). For the arabinose and acetate concentrations determined by HPLC, the average and standard deviation for samples from two individual cultures are presented. Similarly, the WEs were quantitatively analyzed by NMR from two individual cultures. The WE production at different time-points is presented as mg/g CDW to demonstrate the different accumulation patterns of the strains.

Batch cultivations for ADP1 wt and ADP1-ara-aceA were carried out in minimal medium supplemented with 0.1 % casein amino acids, 50 mM Na-acetate and 0.5 % arabinose. Acetate and arabinose concentrations as well as the WE production (at 5, 7, 9 and 10 h time-points) were determined (Figure 5). ADP1 wt consumed all the acetate in approximately 9-10 hours; the amount of biomass increased until the 10-hour time-point, the CDW being 1.3 g/l. The highest amount of WEs was measured at the 9-hour time-point, being 47 mg/g CDW and 60 mg/l. In ADP1 wt, the WEs accounted for 44 % of total lipids. The WE yield was found to be 0.02 g WE/g consumed acetate. The strain ADP1-ara-aceA utilized acetate more steadily and produced less biomass compared to ADP1 wt; the growth ceased after 16 hours along with the arabinose depletion: the biomass remained at 0.6-0.7 g/l CDW between the 16-24 hours of cultivation. Thereafter, the WE content of the cells strongly increased, being highest at 38 h time-point, which also increased the amount of total biomass to 1.0 g/l. The WE titer was found to be 184 mg/l representing 19 % of CDW, which was 3.8-fold higher compared to the ADP1 wt. In addition, the WEs accounted for 80 % of all cellular lipids in ADP1-ara-aceA. The WE yield was 0.08 g WE/g consumed acetate, also being 4-fold higher over the ADP1 wt.

## 4. Discussion

Sugars, mainly glucose, have been the major carbon source for the heterotrophic microbial production of fatty acid derived compounds, such as TAGs and WEs, which can be used for the production of biofuels, biochemicals, and other biocommodities (29, 30). However, in order to increase the feasibility and sustainability of the processes, the possibility to utilize alternative carbon sources is of high interest. Organic acids, such as acetate, serves as a low-cost, abundant carbon source for microbial lipid synthesis. Acetate can be readily derived from the hemicellulose fraction of plant biomass or waste streams, or produced from syngases by microbial fermentation (31). For example, we have previously demonstrated the conversion of carbon dioxide to WEs via acetate intermediate by combining microbial electrosynthesis with aerobic lipid synthesis in a two-stage process (32). Many of the potential acetate streams, however, may have dilute acetate concentrations, thus sustaining conditions that are preferable for biomass production rather than for efficient storage carbon synthesis. On the other hand, highly concentrated acetate feeds can inhibit cell growth (33).

Nitrogen starvation is a commonly used and efficient means to trigger storage lipid accumulation in microbes (34-36). However, in such conditions the cell biomass typically remains low (37), which can result in lower overall product titres. Thus, genetic reprogramming would serve as a means to bypass the natural regulation for storage lipid synthesis. Mechanisms behind the regulation in microbes are not well understood, and therefore it has been challenging to genetically drive cells to overproduce the storage compounds or other fatty acid derived products. Previous strategies include for example the manipulation of the conserved carbon storage regulator CsrA through the CsrA-CsrB ribonucleoprotein complex, by which alterations in the central carbon metabolism and fatty acid synthesis regulation have led to favourable changes in both native and non-native product synthesis pathways in *E. coli* (38). This approach, however, is rather unspecific and potentially difficult to combine with other (targeted) engineering strategies.

In this study, we developed an autonomously regulated circuit for programmable synthesis of WEs in a native production host, *A. baylyi* ADP1. The circuit allows the cells to shift from the biomass mode to the WE synthesis mode independent from the carbon/nitrogen ratio or the growth phase of the culture. In practice, we replaced the native isocitrate lyase aceA with an arabinose inducible system, which allows a conditional and timed knockdown of the expression of *aceA*. This enzyme is essential for the biomass production when the cells grow on acetate. The timed repression of aceA expression is achieved by gradually eliminating the inducer, namely arabinose; the native enzyme activity of glucose dehydrogenase Gcd of *A. baylyi* oxidizes arabinose to arabino-lactone and further to arabonate, which in turn cannot serve as inducers. Importantly, and in contrast to other auto-induction-based systems, arabinose oxidation does not interfere with the utilization of the carbon source, here acetate, and can be thus considered as an orthogonal system. By adjusting the arabinose concentration, a predefined and optimal amount of biomass can be produced. When the inducer concentration is oxidized below the ‘threshold’ concentration, the cells shift from the biomass producing mode to the synthesis mode, efficiently directing carbon to product synthesis.

First, we confirmed that the engineered strain ADP1-ara-aceA with complemented isocitrate lyase was able to grow on acetate as the sole carbon source. We observed that without the presence of arabinose, the cells did not exhibit growth, showing phenotype similar to the knockout strain *A. baylyi* ADP1δ*aceA*∷ *tdk*/*Kan*^*r*^δ*poxB*∷ *Cm*^*r*^. We also observed that arabinose concentration 1% is sufficient to allow the growth of ADP1-ara-aceA to reach at least the same biomass as the ADP1 wt.

Initial arabinose concentrations were in correlation with the obtained biomasses between concentrations 0.2% and 1.0%. The concentration 0.2% was found to be the ‘threshold’ for sufficient biomass production; below this concentration, the cells produced only slightly more biomass compared to the uninduced cells, indicating that arabinose concentrations >0.2% are required for sufficient growth in the studied conditions. This finding was also supported by the reporter induction test; when the cells containing the bacterial luciferase *luxAB* under arabinose promoter were induced with the supernatant from different cultivation time-points (thus having different arabinose concentrations), clear induction of *luxAB* determined as luminescence production was only observed with the sample that contained >0.2% arabinose. According to the semi-quantitative WE analyses, the cultures supplemented with 0.2% or 0.5% arabinose produced the highest WE yields (per biomass). The cultures that were induced with 1% arabinose produced nearly two times more biomass compared to that of the 0.5% culture but had lower WE yield, indicating that a significant proportion of the carbon was directed to the biomass when 1% arabinose was used. Interestingly, the cultures with little (0.1%) or no arabinose produced the lowest WE yield, suggesting that at least subtle levels of *aceA* expression are required, not only for biomass production but also to support WE synthesis. The WE titres (WEs/volume) of 0.5% and 1.0% cultures were estimated to be very close to each other in the two cultures, whereas 0.2% culture had clearly lower volumetric WE production due to low biomass production. Thus, we considered 0.5% arabinose as the most effective inducer concentration in terms of optimal distribution of carbon between biomass and products.

For validation of the system, batch cultures for the ADP1 wt and ADP1-ara-aceA were carried out. It was shown that the engineered strain ADP1-ara-aceA efficiently accumulated WEs in simple batch conditions supplied with relatively low substrate concentration (50 mM acetate) and in non-optimal carbon-nitrogen ratio. The strain ADP1-ara-aceA produced 187 mg/l WEs representing 19 % of the CDW and a yield of 0.08 g WE/g consumed acetate. As expected, the strain grew more slowly and produced less biomass than ADP1 wt, but the yield of WEs per biomass and per consumed acetate were 3.8 and 4 folds higher compared to ADP1 wt, respectively. In addition, the WE titer was found to be 3.1 folds higher compared to that of the ADP1 wt. Thus, the dynamic regulation not only improved the yield of WEs per biomass and per used carbon, but clearly excelled the volumetric titer of that of the wild type strain. For comparison, in a previous study (32), *A. baylyi* ADP1 produced WEs from acetate with a titer of approximately 90 mg/l (from higher initial acetate concentration, 100 mM), with an average yield of 4 % (carbon/carbon), being equal to 0.02 g WE/g consumed acetate. The highest WE titer reported so far has been 450 mg/l, which was obtained when the key enzyme of the pathway (fatty acyl-CoA reductase) was overexpressed and 5 % glucose was used as the substrate (17). However, the yields (0.04g WE/g glucose and 12.5% WEs of CDW) were lower compared to this study. With external alkane supplementation, up to 17% WEs of CDW has been obtained (39).

Notably, the amount of WEs per cell was nearly constant in ADP1 wt at the analysed time points, being 3.9-4.3% of the CDW. In ADP1-ara-aceA, by contrast, the amount of WEs per cell strongly increased along with the arabinose depletion; the percentage of WEs per cell increased from 5.2% to 19% between the sampling points. The highest increase in the WE content (from 7.5 to 19%) was achieved after the arabinose concentration reached the ‘threshold’ 0.2% (equivalent to 15 mM arabinose). Although the shift from biomass mode to lipid mode was rather gradual, the arabinose concentration 0.2% seems to be the key turning point in the cellular state. In the batch culture, initial concentrations 50 mM acetate and 0.5% arabinose were used. By the end of the culture, the arabinose was completely oxidized, and only a small amount (5 mM) of acetate remained unutilized. Thus, by adjusting the substrate and inducer concentrations, the system is potentially scalable to a wide range of substrate concentrations. Moreover, coupling this system with other engineering strategies, such as introducing additional knock-outs (16) and/or overexpression of key enzymes of the pathway (17) could further improve the WE production.

Considering not only the efficient redirection of carbon to product, but also the downstream processing, the purity of the product is important. In ADP1-ara-aceA, WEs constituted 80% of total lipids, indicating high purity of the desired product. In ADP1 wt, only 44% of the total lipids were WEs.

The results from different experiments indicate that at least low levels of isocitrate lyase are required to maintain WE production from acetate, potentially due to the requirements for cells to generate e.g. NADPH for the synthesis. This hypothesis was also supported by a further observation in an additional experimental set-up where the WE content of the *aceA* knockout strain did not increase after a transfer from a glucose medium to an acetate medium (data not shown). While arabinose concentration <0.2% is not sufficient to promote biomass production, it allows the cells to synthetize and maintain the required cofactor balance, and to efficiently produce WEs.

Rapid advancements in the CRISPR/Cas9 technologies have broaden the tools available for targeted genome engineering, and especially the employment of the deactivated Cas9 (dCas9) has recently gained interest in the context of targeted gene silencing (40, 41). While the dCas9 –based tools have been shown to be functional and applicable in a wide range of (microbial) hosts, challenges related to unpredictability, cellular burden and off-targeting may limit its use (42, 43). In addition, at least two different constructs are typically required for the fine-tuned expression of dCas9 and the RNA elements, and the system cannot be easily operated ‘hands-free’ without a timed addition of an inducer. In this context, the system described here provides a straight-forward, readily controllable, and reliable set-up for conditional and timed gene knock-down. In addition, other interesting synthetic biology hosts such as *P. putida* (44, 45) exhibit the same glucose dehydrogenase activity and could thus find this strategy applicable. However, the transferability of this system to others hosts such as *E. coli* and *S. cerevisiae* remains to be investigated in the future.

Our system serves as a simple and scalable method for dynamic, ‘hands-free’ regulation of growth-essential reactions in the cell, which allows targeted and adjusted biomass and product synthesis. Here, the dynamic regulation system was exploited in the conversion of acetate to carbon-rich storage compounds, namely wax esters, that are otherwise not efficiently accumulated in cells without optimized conditions and high carbon-nitrogen ratio. Moreover, the system could be potentially utilized and generalized for a broad range of synthesis pathways that are dependent on acetyl-CoA (17, 46). In addition, other sustainable carbon sources, such as lignin-derived compounds (47-49) would be compatible with this system.

## 5. Conclusions

We showed that an autonomously regulated genetic switch allowed the dynamic decoupling of biomass and wax ester production in engineered *A. baylyi* ADP1, which resulted in 3-4 fold improvements in the wax ester yield and titer compared to the wild type strain. Shifting the cells from a biomass mode to a product synthesis mode was achieved by gradually repressing the growth essential gene *aceA* by a simple and robust set-up. The engineered strain produced 19% WEs of its cell dry weight, being the highest reported among microbes. The study demonstrates the possibility to bypass the challenges related to highly regulated storage lipid synthesis, and strengthens the status of *A. baylyi* ADP1 as a convenient host for metabolic engineering and high-value lipid production from sustainable substrates.

## Acknowledgements

Funding: This work was supported by the Academy of Finland (grant numbers 286450, 310135, 310188, and 311986).

## Abbreviations

WE: wax esters
AceA: isocitrate lyase

## References

1. Tan SZ & Prather KL (2017) Dynamic pathway regulation: recent advances and methods of construction. Curr Opin Chem Biol 41:28–35.

2. Soma Y, Tsuruno K, Wada M, Yokota A, & Hanai T (2014) Metabolic flux redirection from a central metabolic pathway toward a synthetic pathway using a metabolic toggle switch. Metabolic engineering 23:175–184.

3. Soma Y & Hanai T (2015) Self-induced metabolic state switching by a tunable cell density sensor for microbial isopropanol production. Metabolic engineering 30:7–15.

4. Solomon KV, Sanders TM, & Prather KL (2012) A dynamic metabolite valve for the control of central carbon metabolism. Metabolic engineering 14(6):661–671.

5. Brockman IM & Prather KL (2015) Dynamic knockdown of E. coli central metabolism for redirecting fluxes of primary metabolites. Metabolic engineering 28:104–113.

6. Moser F, et al. (2012) Genetic circuit performance under conditions relevant for industrial bioreactors. ACS Synthetic Biology 1(11):555–564.

7. Wältermann M, et al. (2005) Mechanism of lipid-body formation in prokaryotes: how bacteria fatten up. Molecular microbiology 55(3):750–763.

8. Wältermann M & Steinbüchel A (2005) Neutral lipid bodies in prokaryotes: recent insights into structure, formation, and relationship to eukaryotic lipid depots. Journal of bacteriology 187(11):3607–3619.

9. Xu P, Li L, Zhang F, Stephanopoulos G, & Koffas M (2014) Improving fatty acids production by engineering dynamic pathway regulation and metabolic control. Proceedings of the National Academy of Sciences of the United States of America 111(31):11299–11304.

10. Teixeira PG, Ferreira R, Zhou YJ, Siewers V, & Nielsen J (2017) Dynamic regulation of fatty acid pools for improved production of fatty alcohols in Saccharomyces cerevisiae. Microbial Cell Factories 16(1):45.

11. Kurosawa K, Boccazzi P, de Almeida NM, & Sinskey AJ (2010) High-cell-density batch fermentation of Rhodococcus opacus PD630 using a high glucose concentration for triacylglycerol production. J Biotechnol 147(3-4):212– 218.

12. Xu J, Liu N, Qiao K, Vogg S, & Stephanopoulos G (2017) Application of metabolic controls for the maximization of lipid production in semicontinuous fermentation. Proceedings of the National Academy of Sciences of the United States of America 114(27):E5308–E5316.

13. Runguphan W & Keasling JD (2014) Metabolic engineering of Saccharomyces cerevisiae for production of fatty acid-derived biofuels and chemicals. Metabolic engineering 21:103–113.

14. Plassmeier J, Li Y, Rueckert C, & Sinskey AJ (2016) Metabolic engineering Corynebacterium glutamicum to produce triacylglycerols. Metabolic engineering 33:86–97.

15. Tai M & Stephanopoulos G (2013) Engineering the push and pull of lipid biosynthesis in oleaginous yeast Yarrowia lipolytica for biofuel production. Metabolic engineering 15:1–9.

16. Santala S, et al. (2011) Improved triacylglycerol production in *Acinetobacter baylyi* ADP1 by metabolic engineering. Microb Cell Fact 10:36.

17. Lehtinen T, Efimova E, Santala S, & Santala V (2018) Improved fatty aldehyde and wax ester production by overexpression of fatty acyl-CoA reductases. Microbial Cell Factories 17(1):19.

18. Santala S, Efimova E, Koskinen P, Karp MT, & Santala V (2014) Rewiring the wax ester production pathway of *Acinetobacter baylyi* ADP1. ACS Synth Biol 3(3):145–151.

19. Klinke S, Dauner M, Scott G, Kessler B, & Witholt B (2000) Inactivation of isocitrate lyase leads to increased production of medium-chain-length poly(3-hydroxyalkanoates) in Pseudomonas putida. Applied and environmental microbiology 66(3):909–913.

20. Young DM, Parke D, & Ornston LN (2005) Opportunities for genetic investigation afforded by Acinetobacter baylyi, a nutritionally versatile bacterial species that is highly competent for natural transformation. Annual review of microbiology 59:519–551.

21. Kannisto M, Aho T, Karp M, & Santala V (2014) Metabolic engineering of Acinetobacter baylyi ADP1 for improved growth on gluconate and glucose. Applied and environmental microbiology 80(22):7021–7027.

22. Kannisto MS, et al. (2015) Metabolic engineering of Acinetobacter baylyi ADP1 for removal of Clostridium butyricum growth inhibitors produced from lignocellulosic hydrolysates. Biotechnol Biofuels 8(1):1–10.

23. Carr EL, Kampfer P, Patel BK, Gurtler V, & Seviour RJ (2003) Seven novel species of Acinetobacter isolated from activated sludge. International journal of systematic and evolutionary microbiology 53(Pt 4):953–963.

24. Barbe V, et al. (2004) Unique features revealed by the genome sequence of Acinetobacter sp. ADP1, a versatile and naturally transformation competent bacterium. Nucleic acids research 32(19):5766–5779.

25. de Berardinis V, et al. (2008) A complete collection of single-gene deletion mutants of *Acinetobacter baylyi* ADP1. Molecular systems biology 4:174.

26. Santala S, Efimova E, Karp M, & Santala V (2011) Real-time monitoring of intracellular wax ester metabolism. Microb Cell Fact 10:75.

27. Murin CD, Segal K, Bryksin A, & Matsumura I (2012) Expression vectors for *Acinetobacter baylyi* ADP1. Applied and environmental microbiology 78(1):280–283.

28. Santala V, Karp M, & Santala S (2016) Bioluminescence based system for rapid detection of natural transformation. FEMS microbiology letters.

29. Lennen RM & Pfleger BF (2013) Microbial production of fatty acid-derived fuels and chemicals. Current opinion in biotechnology 24(6):1044–1053.

30. Marella ER, Holkenbrink C, Siewers V, & Borodina I (2018) Engineering microbial fatty acid metabolism for biofuels and biochemicals. Current opinion in biotechnology 50:39–46.

31. Bengelsdorf FR, Straub M, & Durre P (2013) Bacterial synthesis gas (syngas) fermentation. Environ Technol 34(13-16):1639–1651.

32. Lehtinen T, et al. (2017) Production of long chain alkyl esters from carbon dioxide and electricity by a two-stage bacterial process. Bioresource Technology 243:30–36.

33. Trček J, Mira NP, & Jarboe LR (2015) Adaptation and tolerance of bacteria against acetic acid. Applied microbiology and biotechnology 99(15):6215–6229.

34. Breuer G, Lamers PP, Martens DE, Draaisma RB, & Wijffels RH (2012) The impact of nitrogen starvation on the dynamics of triacylglycerol accumulation in nine microalgae strains. Bioresource Technology 124:217–226.

35. Alvarez HM & Steinbüchel A (2002) Triacylglycerols in prokaryotic microorganisms. Applied microbiology and biotechnology 60(4):367–376.

36. Ishige T, Tani A, Sakai Y, & Kato N (2003) Wax ester production by bacteria. Current opinion in microbiology 6(3):244–250.

37. Amara S, et al. (2016) Characterization of key triacylglycerol biosynthesis processes in rhodococci. Sci Rep 6:24985.

38. McKee AE, et al. (2012) Manipulation of the carbon storage regulator system for metabolite remodeling and biofuel production in Escherichia coli. Microbial Cell Factories 11:79.

39. Ishige T, et al. (2002) Wax ester production from n-alkanes by Acinetobacter sp. strain M-1: ultrastructure of cellular inclusions and role of acyl coenzyme A reductase. Applied and environmental microbiology 68(3):1192–1195.

40. Larson MH, et al. (2013) CRISPR interference (CRISPRi) for sequence-specific control of gene expression. Nat Protoc 8(11):2180–2196.

41. Peters JM, et al. (2016) A Comprehensive, CRISPR-based Functional Analysis of Essential Genes in Bacteria. Cell 165(6):1493–1506.

42. Nielsen AA & Voigt CA (2014) Multi-input CRISPR/Cas genetic circuits that interface host regulatory networks. Molecular systems biology 10:763.

43. Cui L, et al. (2018) A CRISPRi screen in E. coli reveals sequence-specific toxicity of dCas9. Nat Commun 9(1):1912.

44. Nikel PI & de Lorenzo V (2018) Pseudomonas putida as a functional chassis for industrial biocatalysis: From native biochemistry to trans-metabolism. Metabolic engineering.

45. Nikel PI, Chavarria M, Danchin A, & de Lorenzo V (2016) From dirt to industrial applications: Pseudomonas putida as a Synthetic Biology chassis for hosting harsh biochemical reactions. Curr Opin Chem Biol 34:20–29.

46. Lehtinen T, Santala V, & Santala S (2017) Twin-layer biosensor for real-time monitoring of alkane metabolism. FEMS microbiology letters 364(6).

47. Linger JG, et al. (2014) Lignin valorization through integrated biological funneling and chemical catalysis. Proceedings of the National Academy of Sciences of the United States of America 111(33):12013–12018.

48. Salvachúa D, Karp EM, Nimlos CT, Vardon DR, & Beckham GT (2015) Towards lignin consolidated bioprocessing: simultaneous lignin depolymerization and product generation by bacteria. Green Chemistry doi:10.1039/c5gc01165e.

49. Salmela MM, et al. (2018) Molecular tools for selective recovery and detection of lignin-derived molecules. Green Chemistry.

